# Molecular determinants of Bcl-xL membrane insertion – Structural plasticity exploration in solution and in nanodiscs identifies IsoAsp deamidation as a loss-of function mechanism *in vivo*

**DOI:** 10.64898/2026.05.14.725174

**Authors:** Joann Kervadec, Maria Laura De Sciscio, Adrien Mahler, Mélody Dufossée, Charlène Dupont, Yann Rufin, Axel Boudier-Lemosquet, Chloe James, Stéphane Téletchéa, Stephen Manon, Marco D’Abramo, Muriel Priault

## Abstract

Multi-domain Bcl-2 family proteins share the ability to form dimers and oligomers, regardless of their pro- or anti-apoptotic activity. Homotypic interactions (pro-pro and anti-anti) and heterotypic interactions (pro-anti) are well-documented, but the role of higher-order organization in their survival/death functions and membrane interactions remains largely unresolved. Looking into anti-apoptotic Bcl-xL, essentially engineered/truncated proteoforms lacking the disordered loop and/or the hydrophobic C-terminal helix, have been used as proxies of the full-length (FL) protein, to elaborate on structural transitions and intermediate states between monomers and homooligomers prior to membrane insertion. Using a minimalist approach with recombinant FL-Bcl-xL (aa 1-233) and artificial nano-membranes, we demonstrate that both the loop and the C-terminal helix are potent contributors to Bcl-xL structural plasticity. Unlike 3D domain swapping (3DDS) dimers resolved with the C-terminal truncated protein, FL-Bcl-xL organized in solution as dimers bridging the unique Cys151 from two monomers. This spontaneous fold indicates that the C-terminal helix drives FL-Bcl-xL to explore different conformations than truncated Bcl-xL. Yet, dimerization was not a prerequisite for membrane insertion into nanodiscs and Cys151 did not contribute to Bcl-xL survival functions in cells. These data support monomeric Bcl-xL as the minimal functional unit in membranes. Further exploring the frequently deleted disordered loop, we discovered that deamidation of Asn52 and Asn66 in IsoAsp, but not in Asp, impairs membrane insertion into nanodiscs. Thus, this reductionist biochemical approach clarifies the loss of tumorigenic function we observed for deamidated Bcl-xL in xenograft experiments *in vivo*.

## INTRODUCTION

Apoptosis is a highly regulated form of cell death in metazoans, triggered by diverse stimuli that disrupt cellular homeostasis. A sophisticated network of proteins, in dynamic interactions, keeps a tight check between pro-death and pro-survival events. Among them are the Bcl-2 family proteins, which share structural features (up to four Bcl-2 homology -BH-motifs, and a globular fold essentially composed of α-helices), but are functionally divided into pro- and anti-apoptotic proteins, and a third category of BH3-only regulators. Interplays within this family have been extensively described, notably reporting that these proteins (i) can operate as monomers and as higher order assemblies, (ii) engage either in homotypic and in heterotypic interactions, and (iii) partition between the cytosol and the subcellular membranes (essentially at mitochondria). Importantly, much of the biophysical data has been derived from proteins truncated at their C-terminal hydrophobic helix to overcome solubility issues and accommodate high yields of production, or at their internal disordered loop to reduce sample heterogeneity in structural studies. Therefore, studying unmodified proteins to understand how higher organization influences their survival / death functions and membrane interactions remains a key objective.

Bcl-xL is an anti-apoptotic member of this family. This 26 kDa protein harbors four BH motifs and is organized in nine α-helices. The C-terminal α9 is known to drive membrane anchorage when the protein is bound to mitochondria, while it folds back to occupy the BH3-binding groove when Bcl-xL is cytosolic [1]. A distinctive trait of Bcl-xL is the highly flexible sequence (aa 21-85) described as an intrinsically disordered loop in biophysical assays. This loop is susceptible to post-translational modifications in cellular assays, including phosphorylation, ubiquitylation, proteolytic cleavage and quite uniquely in the Bcl-2 family, the stepwise deamidation of Asn52 and Asn66, which has been investigated in details by our team [2–4].

Structural transitions of Bcl-xL between water-soluble and membrane-bound states have been investigated to address the molecular mechanisms behind its anti-apoptotic function. In this scope, the existence of dimers formed through three-dimensional domain swapping (3DDS) of engineered/truncated proteoforms of Bcl-xL has been discussed as a probable conformational switch required for membrane insertion of Bcl-xL. 3DDS dimers result from the intramolecular rearrangement where α5 and α6 switch from a hairpin to a straightened helix, and where two of these proteoforms intertwine and restore monomeric globular units, with intact BH3 peptide binding pockets [5]. Studies describing the formation of 3DDS dimers indicate detergents [6], heat [7], or alkaline conditions [5] as necessary to induce dimerization. Hence, 3DDS formation is not spontaneous and requires an external trigger to overcome the energetic barrier associated with the structural rearrangement. The analysis of the available crystal structure (PDB: 2B48) ruled out that 3DDS dimers could result from a disulfide bridge engaging the unique Cys151 of two monomers, as these residues are 20.6 Å apart. Finally, 3DDS dimers were described as competent for membrane insertion on the grounds that they displayed higher pore forming activity than monomers [6]. All the above supported the hypothesis that 3DDS dimers might represent the structural transition state preceding Bcl-xL membrane insertion, thus pointing to dimers as the functional unit of anti-apoptotic Bcl-xL in membranes. Yet, this was not consistent with solid state NMR data of Bcl-xLΔloop in nanodiscs, showing that the functional unit in membranes was rather the monomer [1]. Because 3DDS dimers have been observed on C-terminal truncated proteins, we decided to use the genuine un-engineered version of Bcl-xL and investigate *in vitro* its structural plasticity in solution and at the membrane.

Importantly, the treatments (detergents, heat, alkaline solutions) that are conducive to 3DDS dimers, also promote deamidation. This raises the possibility of a mechanistic link between deamidation and dimerization of Bcl-xL, but this issue has inevitably been overlooked when loop-deleted constructs have been considered in the literature. Moreover, deamidation of Bcl-xL has also been shown to narrow the BH3-binding groove and reduce BH3-binding capacity [8,9]. As the same groove accommodates the C-terminal α9 in the soluble conformation of Bcl-xL, an open question is whether deamidation also changes the interactions between α9 and the globular core, thereby influencing the protein membrane partitioning. Together, these observations justify investigating the potential interplay between Asn52/Asn66 deamidation, self-association, and the conformational transitions governing membrane anchorage to advance our understanding of Bcl-xL regulation.

## EXPERIMENTAL PROCEDURES

### Recombinant Proteins

Full-length human Bcl-xL was expressed from the pnYC plasmid [10] in *E. coli* BL21[DE3] cells grown in minimal medium M9 as previously described [3,11]. Briefly, protein expression was induced with 1 mM IPTG (Isopropyl β-D1-thiogalactopyranoside) for 3 h at 37°C. Cells were harvested and lysed in a One Shot cell disruptor in 25 mM Tris-HCl (pH 8.0), 100 mM NaCl, 2 mM MgCl_2_, 10 mM DTT supplemented with 0.1 mg/mL DNase I and protease inhibitors. After centrifugation, the pellet was resuspended in 20 mM Tris-HCl (pH 8.0), 2 mM DTT, 1 mM EDTA, 6 M urea and incubated for 30 min at 4°C under gentle agitation. The insoluble fraction was removed by centrifugation at 30,000 x *g* for 20 min at 4°C, and the supernatant was filtered and loaded onto a size exclusion chromatography column (HighLoad Superdex S-75 16/60, Cytiva) connected to an ÄKTA purifier. The fractions were further submitted to anion exchange chromatography on a HiTrap Q HP column (Cytiva) with a 100 mM to 1 M linear gradient of NaCl. The purified protein was concentrated to 2 mg/mL and aliquoted and stored at -80°C.

### Continuous exchange Cell-Free Protein Synthesis

Full-length human Bcl-xL was cloned between the *NdeI* and *XhoI* sites of the pIVEX 2.3MCS plasmid. A stop codon was introduced to exclude the His_6_-tag as described previously [12]. The C151S mutation was introduced on pIVEX-Bcl-xL using the QuickChange method. Cell-free protein synthesis was performed as described previously [13] in a small volume dialysis chamber (100 μL) containing plasmids and 3.5 µM nanodiscs, separated from a feeding reservoir (1700 μL) by a dialysis membrane (10 kDa MWCO). After protein production, the reaction mix was centrifuged for 15 min at 20,000 x *g* at 4°C to separate the pellet (insoluble proteins) from the supernatant (soluble proteins). The protein content of supernatant and pellet fractions was analyzed by western blotting.

### Production of the Scaffold Protein His7-MSP1E3D1

The protocol to produce and purify the scaffold protein His_7_-MSP1E3D1 was adapted from [13]. Briefly, the plasmid pET28b encoding His_7_-MSP1E3D1 (Addgene) was transformed into *E. coli* BL21[DE3*] and its expression was induced with 1 mM of IPTG for 3 hours. Cells were centrifuged and lysed by sonication. After a centrifugation at 46,000 x *g* for 30 min at 4°C, cell debris were discarded and the supernatant was loaded overnight at 4°C onto a Ni-NTA column (HisTrap FF 1 mL, Cytiva) in a closed circuit. The column was washed successively with 40 mM Tris-HCl (pH 8.0), 300 mM NaCl, 1% Triton X-100 and 40 mM Tris-HCl (pH 8.0), 300 mM NaCl, 50 mM sodium cholate. The column was connected to an ÄKTA purifier and washed with 20 column volume of buffer A (40 mM Tris-HCl (pH 8.0), 300 mM NaCl). The protein was eluted in a two-step process with first, 7% Buffer B (40 mM Tris-HCl (pH 8.0), 300 mM NaCl, 300 mM imidazole) followed by 100% of buffer B. The eluate was dialyzed against a desalting buffer (10 mM Tris-HCl (pH 8.0), 100 mM NaCl), concentrated to 7 mg/mL and stored as working aliquots at -80°C.

### Nanodisc Formation

A mitochondria-like lipid mixture (2.4 mg of POPC, 1.6 mg of POPE, 0.6 mg of DOPS and 0.4 mg of cardiolipin) was assembled in chloroform (Anatrace). Nanodiscs were produced and purification was adapted from [13] using a Superdex 200 Increase 10/300 GL (Cytiva).

### Purification of nanodiscs and associated proteins

Proteins translated in the presence of His-tagged nanodiscs were diluted 1:1 with binding buffer (25 mM Tris (pH 8.0), 250 mM NaCl) and incubated with 80 µL of Ni-NTA Sepharose slurry (Qiagen) for 2 h at 4°C under gentle agitation. The unbound fraction was collected. The resin was washed and bound proteins were eluted in 25 mM Tris-HCl (pH 8.0), 250 mM NaCl, 300 mM imidazole (pH 8.0) in a volume equivalent to that of the unbound fraction. The protein content of unbound and eluate fractions were analyzed by western blot as “nanodiscs-associated proteins”.

### Purification of nanodiscs-inserted proteins

Nanodiscs-associated proteins were further assayed for membrane insertion by sodium carbonate treatment. Briefly, eluates of the previous Ni-NTA affinity chromatography were dialyzed twice for 3 h at 4 °C against binding buffer and then divided into two. For mock treatment, the samples were incubated in binding buffer, whereas the other half was incubated with 100 mM Na_2_CO_3_ (pH 10.0) for 15 min at 4°C. Ni-NTA affinity chromatography was performed again as described above. The protein contents were analyzed by western blot with eluates accounting for “nanodiscs-inserted proteins”.

### Cross-linking assay

Following the purification of nanodiscs and associated proteins, a dialysis was carried out at 4°C against 10 mM HEPES (pH 7.4), 100 mM NaCl. Samples were either mock-treated or incubated with 0.01% Brij 58 at room temperature for 15 minutes to solubilize nanodiscs. Proteins within 7.7 Å were cross-linked with 0.2 mM of disuccinimidylglutarate (DSG) for 30 min at room temperature. The reaction was quenched with 50 mM of Tris-base for 15 min at room temperature. Samples were diluted 1:1 with 2x Laemmli buffer prior to loading onto 4-16% Mini-Protean TGX Precast PAGE gels (Bio-Rad). Adducts in mock conditions represent complexes that were formed in nanodiscs, whereas Brij 58-treated samples reveal complexes that survived solubilization.

### Spontaneous insertion of Bcl-xL into nanodiscs

Recombinant Bcl-xL (2 µg) was either left untreated or subjected to accelerated deamidation with 0.2% NH_4_OH for 16 h at 37°C. Proteins were incubated with 3.5 µM nanodiscs for 1 h at room temperature under gentle agitation. Protein association and protein insertion was assayed as described above.

### Alkaline-induced deamidation

Human recombinant Bcl-xL was stored in 25 mM Tris-HCl (pH 8.0), 450 mM NaCl. For accelerated deamidation, 60 ng of protein was incubated with 0.2% NH_4_OH at 4°C or 37°C for various time periods. The reaction was stopped at -80°C.

### Protein oxidation

Human recombinant Bcl-xL (60 ng) was incubated with 3 mM of CuCl_2_ for 1 h at 4°C. The reaction was stopped by adding 25mM of N-Ethyl-Maleimide (NEM) and 5 mM of EDTA.

### Cell Lines and culture

HeLa (CCL-2) and HCT116 (CCL-247) cells were purchased from the ATCC®. HeLa cells were grown in RPMI and HCT116 cells in McCoy-5A. Media were supplemented 10% fetal calf serum (Dutscher), penicillin (100 U/mL, Invitrogen) and streptomycin (100 µg/mL, Invitrogen). To generate HeLa cells expressing Bcl-xL WT or Bcl-xL C151S, or HCT116 cells overexpressing Bcl-xL WT or Bcl-xLN52DN66A or Bcl-xL N52DN66D, recombinant lentiviruses were engineered, produced and titrated as previously described [14]. The multiplicity of infection was adapted to provide comparable expression levels.

### Animal experiments

This study was conducted in accordance with both Bordeaux University Institutional Committee guidelines (CEEA50) and European Community directives for experimental animal use (L358‐86/609/EEC). Female NMRI-nu mice (age 4-5 weeks) were purchased from Charles River. For xenograft experiments, 1 × 10^6^ HCT116 cells (parental, or overexpressing Bcl-xL WT, monodeamidated Bcl-xL N52DN66A, or doubly deamidated Bcl-xL N52DN66D) were resuspended in 100 µL PBS and injected subcutaneously into the flank of 10 mice per group on day 1. Tumor growth was monitored 3 times a week until a significant differences were observed, for a maximum of 26 days. Tumors were excised and measured.

### Co-immunoprecipitation of Bcl-xL WT or Bcl-xL C151S with Bax WT

HeLa cells were harvested, centrifuged and the pellets were resuspended in 20 mM HEPES (pH 7.5), 150 mM NaCl, 0.1% Triton X-100 and 10% Glycerol). Antibodies against Bcl-xL (rabbit monoclonal anti-human Bcl-xL E18, Abcam) or Bax (mouse monoclonal anti-human Bax 2D2, Santa Cruz Biotechnology) were incubated 1 h at 4°C with magnetic Dynabeads (Invitrogen). Then, 250 µg of total protein extracts was added and incubated under agitation at 4°C overnight. Nonspecific interactions were washed away, and proteins were eluted with 125 mM Tris-HCl (pH 7.0), 4% SDS and 20% glycerol), then heated at 65°C for 5 minutes.

### Apoptosis assays

HeLa cells were were transfected to overexpress Bcl-xL WT or Bcl-xL C151S. Cells were treated with the apoptosis inducer staurosporine at either 200 nM for 16 h or 1 µM for 4 h. Total proteins were extracted in RIPA and 25 µg separated on SDS-PAGE and probed with the indicated antibodies by western blotting.

### Total Proteins Extraction

HeLa cells were harvested and total proteins were extracted in RIPA buffer (1% NP-40, 0.5% sodium-deoxycholate, 0.1% SDS in PBS, supplemented with proteases inhibitor Mini® (Roche Diagnostics)) as described in [4].

### Electrophoresis

Two polyacrylamide gel compositions were used in this study. Standard denaturing Tris/glycine 12.5% SDS-PAGE mini-gels were used with a migration buffer containing 192 mM Glycine, 25 mM Tris and 3.5 mM SDS. Homemade taurine/glycine/imidazole mini-gels described in [4] were formulated to improve the resolution of deamidated species and used with a migration buffer containing 328 mM glycine, 50 mM Tris and 5.2 mM SDS. Electrophoresis was performed at 25 mA per mini-gel.

### Western Blotting

Proteins separated on polyacrylamide gels were transferred onto nitrocellulose membranes (Amersham) saturated with 5% milk in PBS-Tween 20 or TBS-Tween 20. Antibodies used are: rabbit monoclonal anti-Bcl-xL (ab32370, Abcam) 1/4,000 dilution, rabbit monoclonal anti-human Bax (ab182734, Abcam) 1/5,000 dilution, mouse monoclonal anti-his tag (clone 4E3D10H2/E3, Thermo Fisher) 1/5,000 dilution. Horseradish peroxidase-conjugated secondary antibodies were purchased from Jackson ImmunoResearch and used at 1/10,000. Western-blots were developed using Clarity Western ECL (Bio-Rad). Images were captured with a G-Box digital camera (Syngene) and processed using Image J (https://imagej.nih.gov/ij/index.html).

### Computational simulations

The crystal structure of Bcl-xL bound to the Bim BH3 peptide (PDB code: 3FDL) was selected as the input structure for molecular investigations. After removing the ligand, it was used to perform protein-protein docking using HADDOCK 3 web interface (https://github.com/haddocking/haddock3) [15]. The same structure was selected for both monomers, and the sole Cys151 residue in each Bcl-xL monomer was designated as the potential contact region. All other parameters were set to the default. Among the poses obtained, the distance between the sulfur atoms of the Cys151 residue in each monomer was selected as a scoring criterion. The pose with the shortest S-S distance, and characterized by a favorable energetic HADDOCK score, was used as the input for subsequent molecular dynamics (MD) simulations. All the MD simulations were performed using the GROMACS 2024 software package [16] and the CHARMM36 force field [17], as described below. The input structure was centered in a dodecahedral box large enough to avoid periodic conditions, solvated with TIP3P water model and 150 mM NaCl, and the system charge was neutralized. After energy minimization using the steepest descent algorithm, the system was equilibrated in the canonical ensemble by performing a series of short iterative runs, during which the simulation box size was adjusted to reproduce the experimental density of water, as previously described. From the equilibrated structure, six replicates were initiated by assigning different random initial velocities, resulting in an overall time of 11.2 µs. A summary of the simulation time for each run is reported in Table 1.

**Table 1.** Summary of the Bcl-xL non-covalent dimer simulated times, along with the minimum Cys-Cys distance sampled in each replicate.

| Run | Lenght (μs) | Minimum S-S distance (nm) |
| --- | --- | --- |
| 1 | 1.2 | 0.31 |
| 2 | 2.8 | 0.65 |
| 3 | 3.6 | 0.53 |
| 4 | 1.2 | 0.45 |
| 5 | 1.2 | 0.73 |
| 6 | 1.2 | 0.69 |
| Total | 11.2 |  |

An additional simulation of the covalently bound dimer (via a disulfide bridge) was performed using the same procedure described above. The MD-sampled structure with the shortest S-S distance (0.34 nm) was used as the input structure for this additional simulation, after manually creating the S-S covalent bond using PyMOL. All the structural-dynamics analyses were performed using GROMACS, the MDTraj library [18], and in-house Python scripts.

### Statistical analysis

Data are presented as mean ± SEM. Statistical analyses were performed using GraphPad Prism version 8. A value of *p* ≤ 0.05 was considered statistically significant. For *in vitro* studies, data were analyzed using the Mann-Whitney test. For *in vivo* studies, data were analyzed using a repeated-measures mixed-effects model. Geisser–Greenhouse correction was applied to account for potential violations of sphericity. Multiple comparisons between groups were performed using Tukey’s post hoc test with a 95% confidence interval.

## RESULTS

The use of truncated/engineered proteoforms of Bcl-xL has obscured the exploration of its natural structural plasticity in solution and at membranes. To explore the organization of FL-Bcl-xL in solution, we either used Bcl-xL purified from expression in *E. coli* or synthesized by cell-free transcription/translation from bacterial extracts. The former followed the procedure refined by the Marassi group to prevent unwanted bacterial cleavage at Met218 [19], which involves urea denaturation to retrieve Bcl-xL from inclusion bodies. Data obtained from this recombinant purified protein were systematically verified using Bcl-xL transcribed and translated from bacterial extracts, as previously published by our team [12], to avoid focusing on potential artifactual sub-states of the protein possibly induced by urea denaturation.

### FL-Bcl-xL ORGANIZATION IN SOLUTION

To explore Bcl-xL organization in solution, we compared in Figure 1 the effects of conditions previously reported to induce 3DDS dimers formation -heat (Figure 1B), alkaline treatment (Figure 1C) or a combination of both (Figure 1D) – over the time points indicated; as a control, the protein was maintained in storage buffer at 4°C (Figure 1A). The electrophoretic behavior was analyzed on denaturing but non reducing gels. Bcl-xL essentially migrated as a ∼26 kDa monomer at initial time points, but a ∼70 kDa proteoform that could correspond to a dimer, appeared in all conditions. Importantly, the detection of this proteoform in the storage buffer at 4°C indicates that, unlike 3DDS dimers, it forms spontaneously.

**Figure 1:**
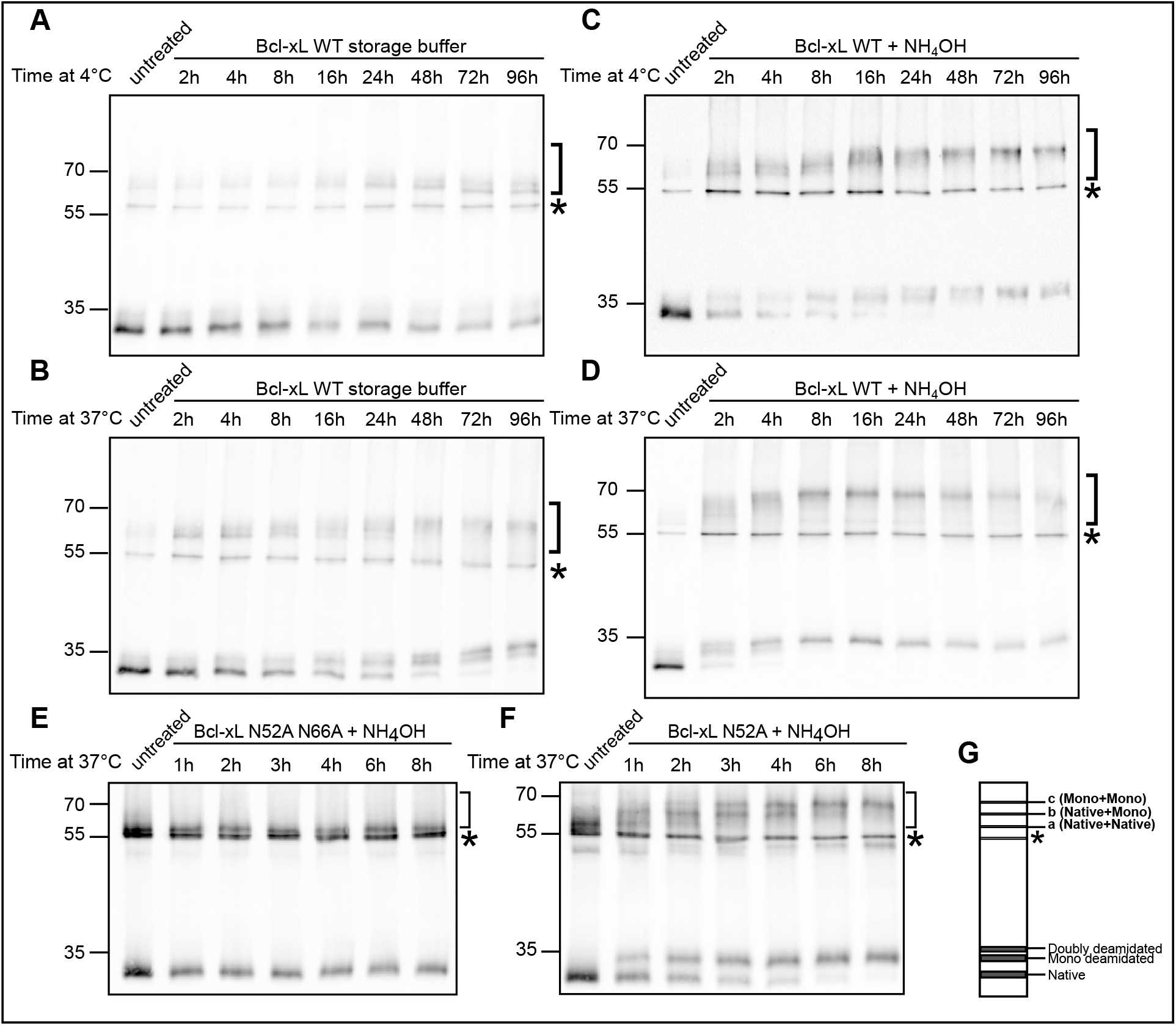
Spontaneous or accelerated deamidation of Bcl-xL. Recombinant Bcl-xL and the indicated mutants were purified from bacteria and incubated for the indicated times and temperatures either in storage buffer (**A and B**) or with 0.2% NH4OH (**C-F**). 10 ng of proteins were separated on 12%TGI gels and analyzed by western blotting against Bcl-xL. Images are representative of at least 8 independent experiments. **G-** schematic representation of Bcl-xL proteoforms separated on TGI gels.] Putative dimers of Bcl-xL, * stacking band at ∼55 kDa characteristic of Bcl-xL separation on TGI gels.

Heat and alkaline treatment are well-established catalysts of deamidation, a spontaneous post-translational reaction in which Asn and Gln residues lose the amino group from their side chain. An intramolecular rearrangement converts these residues into Asp/IsoAsp and Glu/IsoGlu, respectively. We and others have documented that Asn52 and Asn66 in the unstructured loop of Bcl-xL are deamidable [20,21]. Our group further described a Taurine-Glycine-Imidazole (TGI) gel composition suited to use electrophoretic migration distances as a readout of the deamidation status of small proteins, including Bcl-xL [3,4]. TGI gels separate native Bcl-xL as the fastest-migrating band, N52-N66-doubly deamidated Bcl-xL as the slowest migrating band, and monodeamidated species as intermediate (Supplementary Figure 1). As expected, the TGI gels used in Figure 1 allowed monitoring time-dependent migration shifts of monomeric Bcl-xL exposed to heat or ammonia, with significant acceleration when both were combined. Strikingly, the migration shifts observed for the monomeric protein were strictly mirrored by the ∼70 kDa proteoform. This raised the question of whether deamidation and dimerization are interdependent, a matter overlooked in all the studies where the intrinsically disordered region (IDR) of Bcl-xL was deleted. We reasoned that if deamidation were required for dimerization, a non-deamidable mutant (N52A/N66A) would prevent formation of the ∼70 kDa proteoform. Figure 1E shows that heat+ammonia-exposed Bcl-xLN52AN66A did not exhibit any migration shift of the monomer over time, as expected. However, the ∼70 kDa proteoform was still detected, indicating that deamidation is not a prerequisite for dimerization. To examine if dimers can form regardless of the monomer deamidation status, we used a single mutant where only N66 is deamidable (mutant N52A in Figure 1F). We reasoned that if dimers were homotypic, N52A would only dimerize with itself, forming band “a” in Figure 1G; then, as N66 deamidates, homotypic dimers would consist solely of deamidated (band “c”). The observation of an intermediate band “b” indicates that heterotypic dimers are also present. Finally, band “a” being consumed over time in Figure 1F indicates that Asn deamidation occurs not only in Bcl-xL as a monomer, but also in dimers. This set of experiments provides unprecedented evidence that Bcl-xL deamidation and dimerization proceed independently.

### FL-Bcl-xL FORMS SPONTANEOUS OXIDATIVE DIMERS IN SOLUTION

When we analyzed the electrophoretic mobility of recombinant Bcl-xL samples from Figure 1 in the presence of β-mercaptoethanol (β-ME) as a reducing agent (Figure 2B), or the Bcl-xL C151S mutant (Figure 2C), the ∼70 kDa proteoform was no longer detected compared to untreated samples (Figure 2A). Conversely, enforced oxidation with CuCl_2_ increased dimerization of WT Bcl-xL but not of Bcl-xL C151S (Figure 2D). These results demonstrate that the ∼70kDa proteoform indeed resulted from a disulfide bridge between the unique Cys151 residues in helix α5 of FL-Bcl-xL monomers. This finding confirms that the proteoform studied here differs from 3DDS dimers, in which the two Cys151 are too far apart (20.6 Å) to form a disulfide bridge. Therefore, FL-Bcl-xL in solution spontaneously organizes not only as monomers but also shows intrinsic flexibility conducive to a dimeric substate where Cys151 becomes solvent-exposed, whereas it is otherwise shielded in the monomer structure (PDB 1LXL) (Figure 2E).

**Figure 2:**
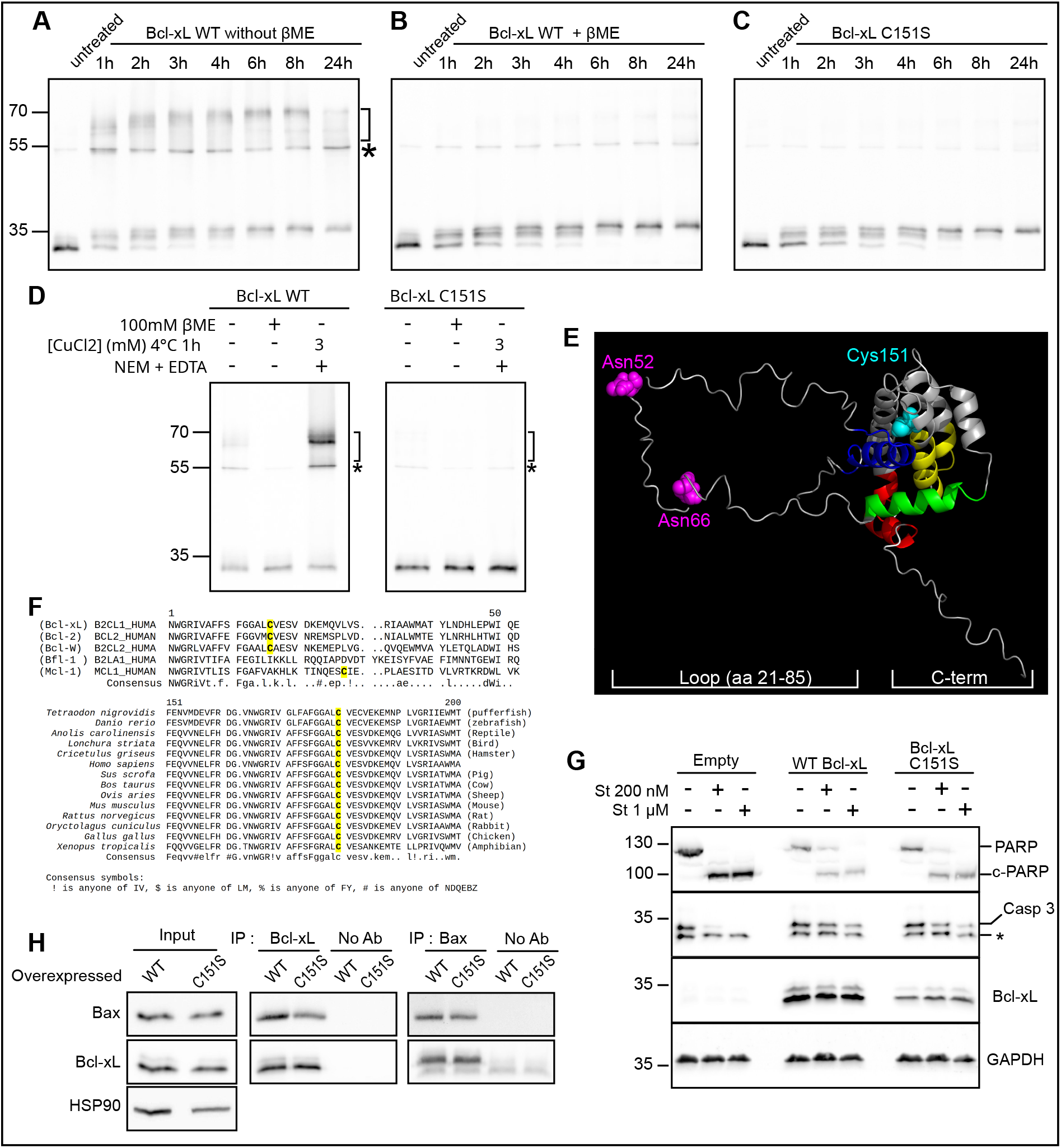
Bcl-xL C151-dependent dimers: **A-C** Recombinant Bcl-xL WT or Bcl-xL C151S were purified from bacteria and incubated at 37°C with 0.2% NH4OH for the indicated times. 10 ng of proteins with or without β-mercaptoethanol were separated on 12%TGI gels. Images are representative of at least 4 independent experiments. **D-** Recombinant Bcl-xL WT or Bcl-xL C151S were mixed with sample buffer with or without β-mercaptoethanol, or they were incubated for 1 h in oxidizing conditions with CuCl_2_. The reaction was stopped by 5 mM EDTA and the remaining reactive cysteines were alkylated with 25 mM NEM. Images are representative of at least 5 independent experiments. **E-** NMR structure of Bcl-xL (PBD: 1LXL) highligthing C151, deamidable N52 and 66, and the domains of the protein frequently deleted in biophysical studies. **F-** Sequence alignments showing conservation of C151 in other human anti-apoptotic members, and in Bcl-xL across metazoan species. **G-** HeLa cells overexpressing WT Bcl-xL or Bcl-xL C151S were treated with staurosporine at 200 nM for 16 h, or at 1 µM for 4 h. Total proteins were extracted and 25 µg separated on SDS-PAGE. Western blots showing Bcl-xL, PARP, cleaved c-PARP, Caspase 3 and GAPDH are representative of at least 3 independent experiments. **H-** HeLa cells overexpressing the indicated proteins were lysed in a Triton X-100 buffer and 250 µg of total proteins were submitted to immunoprecipitations by antibodies to Bcl-xL or Bax, or without antibodies for unspecific detection. Proteins eluted were separated on 12% SDS PAGE followed by immunodetection. HSP90 was used as a loading control. * unspecific band,] redox dimers.

To explore the possibility of the formation of disulfide bridge-mediated dimers, we performed computational simulations. Experimental structures of FL-Bcl-xL, as a starting point, are not available, however NMR analysis have determined that intramolecular interaction of α9 with the BH3-binding groove is very similar to Bcl-xL association with BH3 peptides [1]. Therefore, we used the X-ray structure of the Bcl-xL monomer bound to Bim BH3 peptide (PDB code: 3FDL) as the starting point for docking calculations, designating Cys151 as the reactive site in both monomers. The best pose obtained was compatible with a Cys151-Cys151 bond between the monomers: the residues, located at the center of the complex, were 1.1 nm apart, and geometric / electrostatic analyses revealed no steric or charge clashes, highlighting instead strong complementarity (Figure S2). We further analyzed the conformational dynamics of this non-covalent dimer by performing six molecular dynamics (MD) simulations in water with 150 mM NaCl, starting from the best docking pose (see Methods). In four of the six replicates, the initial 1.1 nm distance between the two Cys151 residues was maintained throughout the simulation (Figure 3A), indicating that the complex remained stable. Remarkably, in one replicate (run1), a transition occurred after ∼300 ns (Figure 3B), resulting in a more compact structure in which the central helices were brought closer together, and the two Cys151 residues reached a minimum distance of 0.4 nm (Figure 3C-D); this new conformation remained stable for the rest of the simulation. A similar event also occurred in another replicate (run4), although reaching a slightly different conformational basin characterized by a greater average Cys151-Cys151 distance than in run1. Notably, even in replicates where no conformational transition was observed, the minimum Cys151-Cys151 distance sampled was consistently below 0.8 nm (Table 1), a value significantly lower than in the crystallographic structure of the 3DDS dimers. Together, these computational simulations support the experimental hypothesis that Cys151-Cys151 dimers form spontaneously in solution. However, the kinetics of experimental dimerization may be much slower than the MD simulations time scale; indeed, extending the simulation time to over 2.5 µs for run2 and run3, did not reveal any transitions (Figure S3). Further analysis of the structural dynamics showed that helices α1 exhibits the highest mobility relative to other regions of the non-covalent dimer (Figures S3 and S5). RMSD fluctuations indicate that both the position and orientation of helices α1 deviate significantly from the docked structure (Figure S3B, S3C and S3D). Conversely, the two pairs of central helices α5 and α6 are more stable across replicates ; in Figure S3E, the increase in RMSD values for run1 and run4 reflects the sliding movement of α5 and α6 depicted in Figure 3, which reduces the distance between the two Cys151. Remarkably, this sliding coincides with the concomitant formation of inter-monomer salt bridges and hydrogen bonds involving selected residues (Figure S4), further supporting the strong directional interactions between monomers. These interactions stabilize a complex protein complex formation in which the two Cys151 are the closest (d < 0.7 nm).

**Figure 3:**
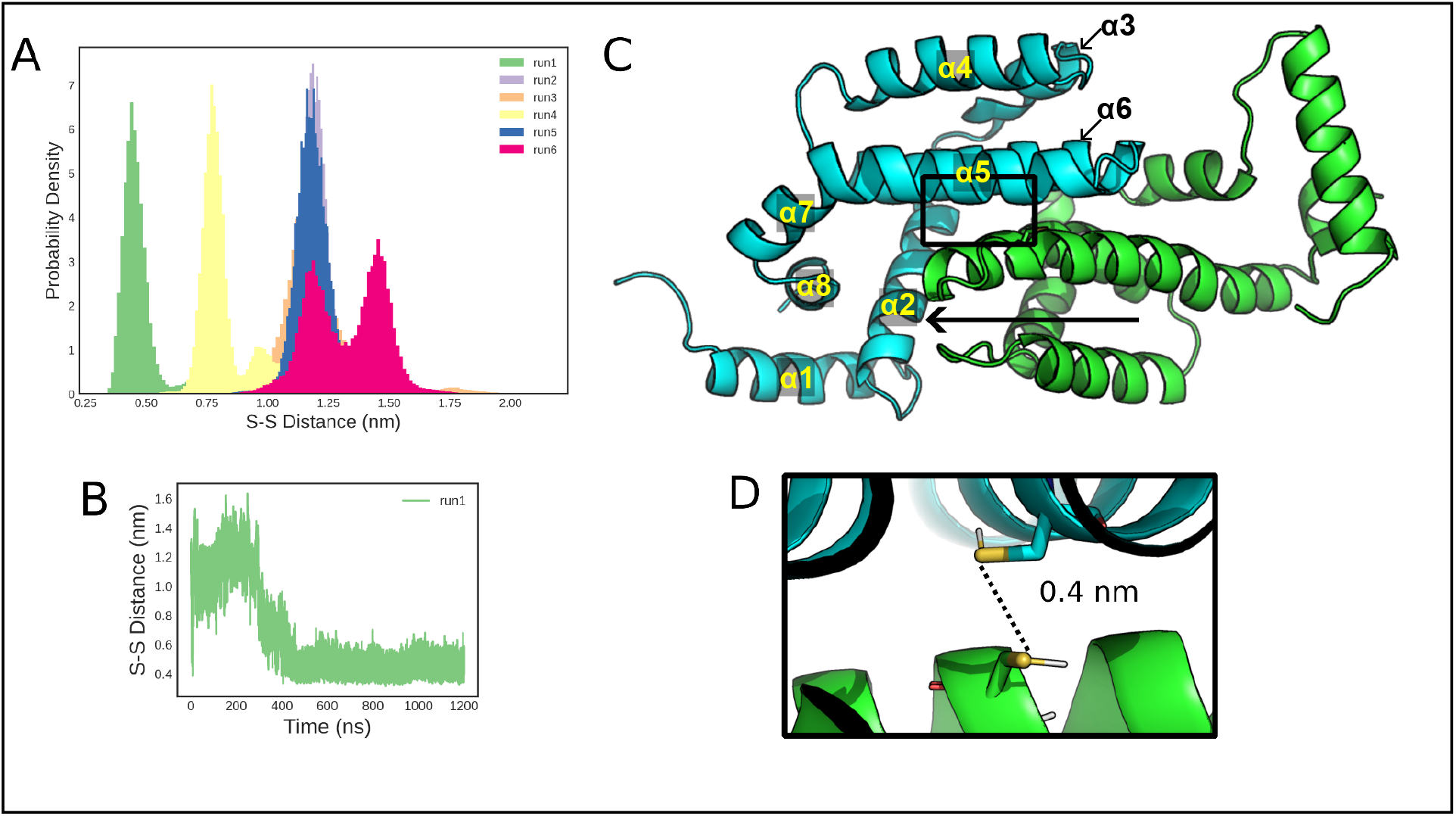
Spontaneous conformation rearrangement of Bcl-xL non-covalent dimer, potentially enabling inter-monomer disulfide bridge formation. **A-** Probability density of the Euclidean distance between the two sulfur atoms of each monomer’s Cys151 sidechain (S-S distance), for all the MD replicates. Data indicate that the distance between the two Cys151 varies depending on the simulations run on unbound monomers. **B-** Temporal evolution of the S-S distance for run1 in A: data show that the structural transition bringing both Cys151 the closest occurs at ∼300 ns of simulation, and that the compact structure remains stable over time. **C-D-** Bcl-xL non-covalent dimer structure resulting in the minimum S-S distance sampled along the MD simulations. Each monomer, in cartoon representation, is colored either cyan or green. The arrow indicates the sliding movement of the central hairpin helices α5 and α6, allowing to shorten the S-S distance. The D inset shows a zoomed-in view of the two Cys151 residues in a conformation compatible with the formation of a disulfide bridge.

The experimental data above suggest that the C-terminal hydrophobic domain of Bcl-xL favors a dimerization pathway distinct from 3DDS in solution, and that this fold forms spontaneously, without overcoming any energetic barrier. Cys151 is located at the end of helix α5, which is highly conserved among Bcl-2 family members in *Homo sapiens* and across metazoan species (Figure 2F). Given that sequence conservation is often indicative of a conserved function, we investigated the role of Cys151 in Bcl-xL-mediated apoptosis regulation.

The pan-kinase inhibitor staurosporine was used to challenge the viability of HeLa cells over-expressing either WT Bcl-xL or Bcl-xL C151S. Cleavage of PARP and Caspase 3 was delayed as efficiently by WT Bcl-xL and Bcl-xL C151S (Figure 2G). Similarly, Bax interacted as efficiently with both proteins (Figure 2H). Additionally, the C151S mutation did not modify Bcl-xL autophagic function in serum-deprived HeLa cells (data not shown). Thus, preventing the formation of Cys151-dependent Bcl-xL dimers does not modify cell survival regulation, at least in assays involving differentiated cell lines. Experiments involving enforced oxidation of cellular extracts failed to generate a 70 kDa proteoform (data not shown), suggesting that redox dimers may not form in cells. However, our previous experiments focused on the organization of FL-Bcl-xL in solution, while it has been long established that, in cells, Bcl-xL predominantly localizes to membranes (mitochondria and ER) [22,23]. Therefore it was necessary to explore how recombinant FL-Bcl-xL partitions to membranes, to determine if Cys151-dependent dimers represent a preliminary sub-state prior to membrane insertion.

### FL-Bcl-xL ORGANIZATION IN NANODISCS

Nanodiscs (ND) are stable synthetic nano-membranes stabilized by apolipoprotein-derived membrane-scaffolding proteins (MSP), which have significantly advanced the biochemical and structural study of membrane proteins [24]. Our team has successfully used cell-free protein synthesis in the presence of preformed NDs to couple translation with membrane-insertion of Bcl-2 proteins [13,25] including Bcl-xL [12]. In particular, we established that this system is well suited to study how Bcl-xL partitions between solution and NDs. Figure 4A presents the flowchart in which Bcl-xL WT and Bcl-xL C151S are synthesized in the presence of NDs stabilized by HIS_7_-MSP1E3D1. A first His-Trap affinity chromatography step separates Bcl-xL that co-elutes with NDs, from soluble (unbound) Bcl-xL. Comparing Bcl-xL WT and Bcl-xL C151S shows that both proteins are equally competent for membrane association (quantified in Figure 4B). Membrane association is further distinguished from membrane insertion by treating the eluate fraction of His-Trap #1 with sodium carbonate to strip loosely associated Bcl-xL, followed by a second His-Trap (#2). Only ND-inserted Bcl-xL is recovered in the eluate from this second chromatography step. Comparing Bcl-xL WT and Bcl-xL C151S indicates, this time, that both proteins are equally competent for membrane insertion (quantified in Figure 4C). To further examine if dimers or higher order assemblies are stabilized after insertion into NDs, eluates from His-Trap #2 were treated with Brij 58, a non ionic surfactant used at ∼10x CMC to disassembles NDs, followed by cross-linking with disuccinimidyl-glutarate (DSG), an amine-reactive homobifunctional crosslinker. SDS-PAGE separation of Brij 58-treated and untreated samples revealed Bcl-xL-containing adducts formed in NDs that persisted despite ND destabilization by the surfactant. In contrast, separation of untreated samples revealed all proteoform assemblies within the 7.7 Å DGS. These include Bcl-xL-(n) adducts as well as Bcl-xL(x)-MSP1E3D1(y) adducts. Figure 4D shows that the same adducts are formed by Bcl-xL WT and Bcl-xL C151S. Together, these results indicate that preventing the formation of Cys151-dependent dimers has no effect on the partitioning of Bcl-xL between the soluble phase and membranes, or on its supramolecular assembly in NDs. This suggests that the Cys151-driven dimeric proteoform is not a mandatory intermediate prior to membrane insertion. These experiments argue in favor of monomeric Bcl-xL as the minimal functional unit for translocation to membranes.

**Figure 4:**
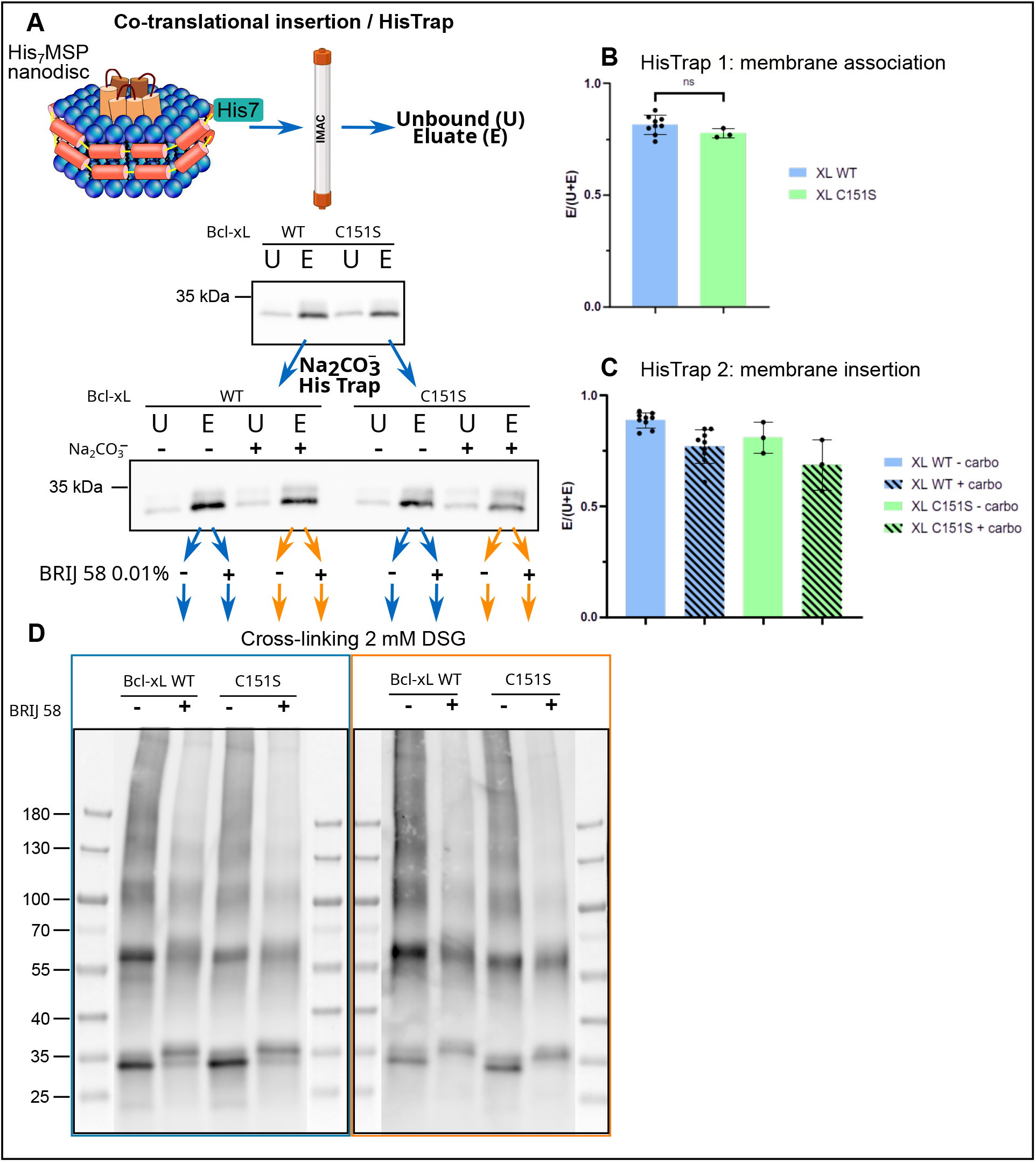
Preventing C151-driven Bcl-xL dimers does not alter membrane interaction nor supramolecular organization. **A-** Bcl-xL WT or Bcl-xL C151S were produced from cell-free bacterial extracts in the presence of preformed nanodiscs (NDs) stabilized by His_7_-MSP1E3D1. His-Trap #1 separates soluble Bcl-xL (unbound) from ND-associated Bcl-xL (Eluate). Fractions were separated by SDS-PAGE and immunodetection of Bcl-xL performed. **B-** Densitometric analyses of n>3 indepedent experiments were quantified. **C-** Eluates from His-Trap #1 were treated with sodium carbonate and further submitted to His-Trap #2 to separate Bcl-xL loosely bound to NDs (unbound) from ND-inserted Bcl-xL (Eluate). Fractions were separated by SDS-PAGE and immunodetection of Bcl-xL was performed. Densitometric analyses of n>3 independent experiments were quantified. **D-** Eluate fractions from His-Trap #2 were incubated with 0.01% Brij 58 (15 min, RT) to disassemble NDs, followed by 2 mM DSG (disuccinimidylglutarate) cross-linking (30 min, RT). Samples were separated on denaturing gradient 4-16% TGX Precast PAGE, and immunodetection of Bcl-xL performed.

### INFLUENCE OF Bcl-xL INTRINSICALLY DISORDERED REGION IN PARTITION TO MEMBRANES

Having shown that the presence of the C-terminal hydrophobic helix drives FL-Bcl-xL to adopt a dimeric conformation distinct from that of Bcl-xLΔC-ter, we aimed to complete the exploration of commonly deleted regions of the protein by focusing on the role played by the IDR in Bcl-xL partitioning to membrane.

Previous studies have reported that post-translational modifications (PTMs) in the IDR result in intramolecular interactions regulating the geometry of the Bcl-xL surface groove which binds BH3-containing partners. This same surface grove is predicted to accommodate the C-terminal hydrophobic helix of unbound Bcl-xL in solution [26,27]. Hence, PTMs of the IDR may facilitate the structural transition between soluble and membrane-anchored Bcl-xL by promoting extrusion of the terminal helix.

Particularly relevant to the present study, deamidation in the IDR has been proposed to alter the loop dynamics, inducing allosteric narrowing of the BH3-binding grove [8,9]. The observation that Bcl-xL deamidation is inhibited in primary cells from patients with hematological malignancies led to consider that this PTM entails a loss of anti-apoptotic functions [28]. However, cellular assays using Asp deamido-mimics failed to clarify the underlying mechanism: compared to Bcl-xL WT, these mutants exhibited unchanged proteostasis, subcellular localization, ability to oppose apoptosis and interactions with pro-apoptotic partners [20]. Further evidence was therefore needed to confirm that deamidation inhibits Bcl-xL tumorogenic function *in vivo*. We performed xenograft experiments in nude mice: even though statistical significance was not reached, an obvious trend indicated that tumors formed from HCT116 cells overexpressing Bcl-xL were larger than those from parental HCT116 cells (Figure 5A), as expected for an oncogene, and consistent with the literature [29]. In contrast, tumors from cells overexpressing Asp deamido-mimics of Bcl-xL (mono- or doubly-deamidated) were indistinguishable from those of parental HCT116 cells. Importantly, these results extend to solid tumors the observation that Bcl-xL deamidation limits tumorigenicity *in vivo*.

**Figure 5:**
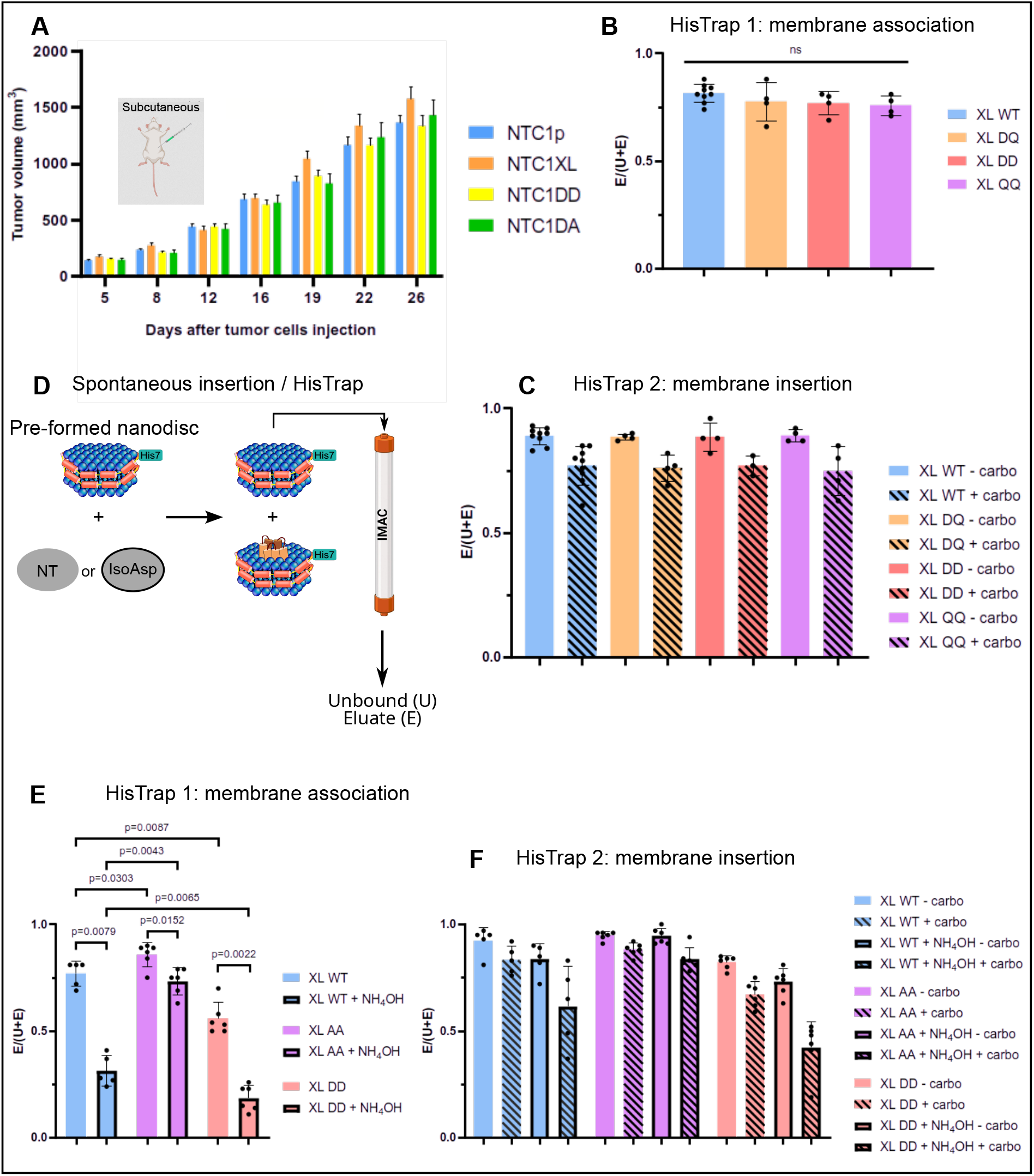
Iso-Asp role in deamidation-driven loss of function of Bcl-xL. **A-** HCT116 cells transduced to overexpress similar amount of Bcl-xL WT (NTC1XL) or the Asp-deamidomimics in position 52 only (NTC1DA) or 52 and 66 (NTC1DD) were xenografted in nude mice, and tumor growth followed for the indicated time, compared to parental cells (NTC1p). **B-** Proteins produced from cell-free bacterial extracts in the presence of nanodiscs (NDs) stabilized by His_7_-MSP1E3D1 were submitted to His-Trap #1 to separate soluble proteins (Unbound) from ND-associated proteins (Eluate). Fractions were separated by SDS-PAGE and immunodetection of Bcl-xL performed. Densitometric analyses of n>4 independent experiments were quantified. **C-** Eluates from His-Trap #1 were treated with sodium carbonate and further submitted to His-Trap #2 to separate proteins loosely bound to NDs (Unbound) from ND-inserted proteins (Eluate). Fractions were separated by SDS-PAGE and imunodetection of Bcl-xL performed. Densitometric analyses of n>3 independent experiments were quantified. **D-** Flowchart of the assay for spontaneous insertion of untreated (NT) or IsoAsp deamidated recombinant proteins in pre-formed NDs. **E-** Recombinant Bcl-xL or the indicated mutants, treated or not with 0.2% NH_4_OH to produce IsoAsp deamidation were incubated in the presence of preformed NDs. Associated proteins were assayed as in B (n>5). **F-** Membrane-inserted proteins were assayed as in C (n>5). AA = Bcl-xL N52A N66A; DD = Bcl-xL N52D N66D.

We then transitioned to the reductionist nanodisc-based assay to quantitatively compare the co-translational membrane association and insertion of Bcl-xL variants differing in deamidation state. To recapitulate deamidation, positions 52 and/or 66 were substituted with Asp; conversely, the non-deamidable mutant was generated by substituting residues 52 and 66 with Gln to preserve a polarity comparable to Asn, while minimizing spontaneous deamidation. Figure 5B shows that all the proteins exhibited similar ND-association, and carbonate treatment revealed that comparable insertion into NDs was achieved, regardless of the proteins deamidation state (Figure 5C). Hence, this assay further reinforced the apparent paradox between i*n vivo* and *in vitro* findings.

Following the rationale that Asp residues in proteins are prone to isomerization [30,31], we hypothesized that IsoAsp, rather than Asp, might account for the loss of function *in vivo*. To test this, we exploited the unique advantage provided by *in vitro* studies to compare Bcl-xL harboring either Asp or IsoAsp at positions 52 and 66. Indeed, IsoAsp cannot be encoded by DNA for incorporation into *de novo* synthesized proteins, but it is the predominant species (∼85% IsoAsp vs ∼15% Asp) produced upon opening of the succinimide cyclic intermediate during accelerated deamidation by alkaline treatments like NH_4_OH [32,33]. Preformed nanodiscs were this time used to compare the spontaneous association/insertion of recombinant Full-Length proteins purified as previously described [4] and treated or not with 0.2% NH_4_OH. These experiments demonstrated that Bcl-xL treated with ammonia significantly lost its ability to associate with NDs. Importantly, this effect could not be attributed to nonspecific structural damage following alkaline treatment, as we previously confirmed by circular dichroism that the protein fold is retained [3]. This possibility was further ruled out by the association rate observed with the non-deamidable protein (Bcl-xL AA) exposed to the same alkaline treatment. Interestingly, the deamido-mimic mutant Bcl-xL DD exhibited behavior similar to that of the WT protein (Figure 5E).

Finally, Figure 5F shows that IsoAsp not only impaired association, but also insertion of Bcl-xL into NDs, as observed after carbonate treatment. Together, our findings provide an explanation to the discrepancy between the loss of function of deamidated *in vivo* as opposed to the absence of phenotype *in cellulo*. Additional work will be required to determine if, indeed, IsoAsp deamidation prevails over Asp deamidation *in vivo*, and to explain why it does not in cultured cells.

## DISCUSSION

Homotypic and heterotypic interactions are features shared by Bcl-2 family proteins, which have long been studied both for the pro-apoptotic multidomain proteins Bax and Bak, and for the anti-apoptotic members Bcl-2, Bcl-xL and, more recently, Bcl-w [34]. To investigate how these interactions influence in the cytosol-to-membrane partitioning of Bcl-2 family members, truncated proteins and/or point mutants have repeatedly been used, the former to facilitate purification and reduce sample heterogeneity, and the latter to capture intermediate conformations. However, recent technological developments now enable studies using full-length proteins, which will provide the indispensable information to validate or refute the current models under debate.

Our choice to use full-length Bcl-xL in the present study revealed that the protein exhibits substantial intrinsic plasticity in the absence of detergents. The spontaneous formation of Cys151-dependent dimers (Figure 1 and 2) suggests that the C-terminal hydrophobic helix guides structural trajectories for the protein in solution that differ from those, like the 3DDS structure, previously documented for engineered/truncated Bcl-xL; this observation further emphasizes that truncated constructs may adopt unrealistic conformations, and thus, systematic extrapolation to the intact protein should not be inferred. Our computational analyses support that Bcl-xL explores a broad conformational landscape (Figure 3). Indeed, the simulations initiated with two monomers generated various models of distinct yet consistently stable structures; Cys151 is located in helix α5, which composes a central hairpin with α6. By means of extended MD simulations, we documented a directional sliding of these hairpins (Figure S4) bringing the monomers closer together, resulting in a more compact structure. Inter-monomer salt bridges and hydrogen bonds involving various residues stabilize this structure, in which inter-molecular distances between the two Cys151 are compatible with the formation of redox dimer. Of note, computational simulations could not determine whether Cys151-linked dimers retain the ability to engage BH3-containing partners. Several residues critical for BH3 recognition are located in immediate spatial proximity to Cys151, including residues between helices α2 and α3 (Lys99, Arg100, Tyr101), Leu108 in α3, and residues in α4 (Val126, Glu129, Arg132) and α5 (Gly138, Arg139, Val141, Phe146) [35]. Because Cys151 lies adjacent to this functional interface, the formation of a disulfide bridge may very well induce local rearrangements that affect the geometry and accessibility of the BH3-binding groove. Unfortunately, a high-resolution structure of FL-Bcl-xL bound to FL-BH3 partners has yet to be determined, and the partial structures available with BH3-peptides cannot capture the steric constraints imposed by complete binding interfaces between whole proteins. Therefore, insufficient data are currently available to confidently assess, through molecular dynamics simulations alone, BH3-binding capacity of Cys151-dependent dimers.

Whether Cys151-dependent dimers represent a biological organization remains an open question. Indeed, the absence of detectable redox dimers in our cellular studies (Figure 2) does not support these assemblies as dominant functional species. However, the conservation of Cys151 across Bcl-2 family members and metazoans conversely suggests a selective advantage. This paradox raises several possibilities: (i) that Cys151-mediated dimerization might play a role in non-apoptotic processes, such as Ca^2+^ signaling or mitochondrial dynamics, that depend on cellular context or binding partners not tested here ; (ii) alternatively, the evolutionary pressure maintaining this residue may operate during developmental stages where oxidative stress and protein homeostasis differ from those in differentiated cell lines; (iii) lastly, the absence of redox dimers in cells may reflect an active counter-selection process to avoid this fold and instead stabilize the monomer. Our *in vitro* experiments highlight the spontaneous propensity of Bcl-xL to adopt a disulfide-linked conformation, but indicate that dimerization is not a prerequisite for membrane insertion (Figure 4). In cells, it is possible that chaperoning mechanisms or binding to partners stabilize productive conformations and prevent these assemblies if they are detrimental to apoptosis regulation. In this respect, rather than suggesting that redox dimers are artifactual *in vitro* conformations, our findings could reveal transient folds that, if stabilized in cells, could lead to nonfunctional assemblies. This structural regulation could represent a new avenue for anti-cancer strategies through restriction of Bcl-xL to nonproductive conformations.

Within the Bcl-2 family, Bcl-xL is additionally singular due to its unique susceptibility to spontaneous deamidation, raising the possibility of a mechanistic connection between the natural instability of Asn52 and Asn66, the protein’s ability to self-associate, and its structural plasticity to transition from a soluble to a membrane-anchored state. To our knowledge, whether deamidation contributes to homotypic interactions, or conversely whether homotypic interactions modulate the kinetics of Bcl-xL deamidation, has not been investigated, nor how these processes influence Bcl-xL membrane anchorage. The present study established that Bcl-xL homotypic interactions and deamidation are not interdependent, but proceed as parallel pathways (Figure 1). This work also provides an unprecedented exploration of the existing discrepancy between deamidated Bcl-xL loss of anti-apoptotic function *in vivo* and the lack of phenotype *in cellulo*. Our experiments extended to solid tumors the anti-tumorogenic effect of deamidated Bcl-xL, previously inferred from observations that Bcl-xL deamidation is inhibited in myeloproliferative disorders [28]. Yet, implementing a nanodiscs-based reductionist approach to dissect the mechanism at play, we found that, consistent with cellular assays [20], Asp deamido-mimics are indistinguishable from the native protein in terms of membrane association or insertion (Figure 5). Importantly, however, we discovered a functional divergence between the proteoforms produced by deamidation: our data revealed that only IsoAsp deamidation impairs Bcl-xL association with and insertion into nanodiscs. Further studies are needed to determine whether IsoAsp deamidation is predominant *in vivo*, and why this would not occur in cultured cells. In particular, it will be useful to assay the activity of the repair enzyme PIMT or the cells content in S-adenosine methionine, as both are critical to control the level of IsoAsp-containing proteins [36].

A previous study by Follis et al. using solid state NMR of Bcl-xL embedded in nanodiscs, demonstrated that the introduction of a negative charge within the flexible loop, either through phosphorylation or deamidation, promotes intramolecular interactions between the loop and a positively charged Arg cluster on the globular core [8]. In turn, an allosteric regulation narrows the BH3-binding grove and decreases affinity for BH3-containing ligands. However, this study did not distinguish between Asp and IsoAsp deamidation products, nor did it address the dynamics of Bcl-xL association to membranes. Our data indicate that both are critical parts of the mechanistic aspect of Bcl-xL regulation by deamidation. Although Asp and IsoAsp generate identical net charges, they differ profoundly in geometry: IsoAsp inserts an additional methylene group into the peptide backbone, thereby generating additional structural distortions. Our findings therefore suggest that the biological consequences of Bcl-xL deamidation are not solely driven by charge redistribution within the loop, but also by conformational remodeling that selectively interferes with membrane insertion.

Using full-length Bcl-xL, this study advanced knowledge by providing novel structural and functional insights. It uncovered the role of the C-terminal helix in exploring conformational space distinct from truncated proteoforms; the spontaneous formation of Cys151-dependent dimers *in vitro* raises the question of their absence *in cellulo* and opens innovative hypotheses about unknown chaperoning processes. Our findings related to the unstructured loop also allowed to progress in solving the paradox of deamidated Bcl-xL loss of function *in vivo* by showing that IsoAsp but not Asp deamidation selectively impairs membrane insertion.

## ACKNOWLEDGMENTS

This work was supported by Agence Nationale de la Recherche ANR-22-CE17-0050 to MP, CJ and ST. This work was also supported by Ligue Nationale contre le Cancer to MP.

We are thankful to Sophie North-Chassande and Benoît Rousseau (Université de Bordeaux) for their help with animal experiments.

## Notes

### Competing Interest Statement

The authors have declared no competing interest.

