## supplementary data for "Molecular determinants of Bcl-xL membrane insertion – Structural plasticity exploration in solution and in nanodiscs identifies IsoAsp deamidation as a loss-of function mechanism *in vivo*"

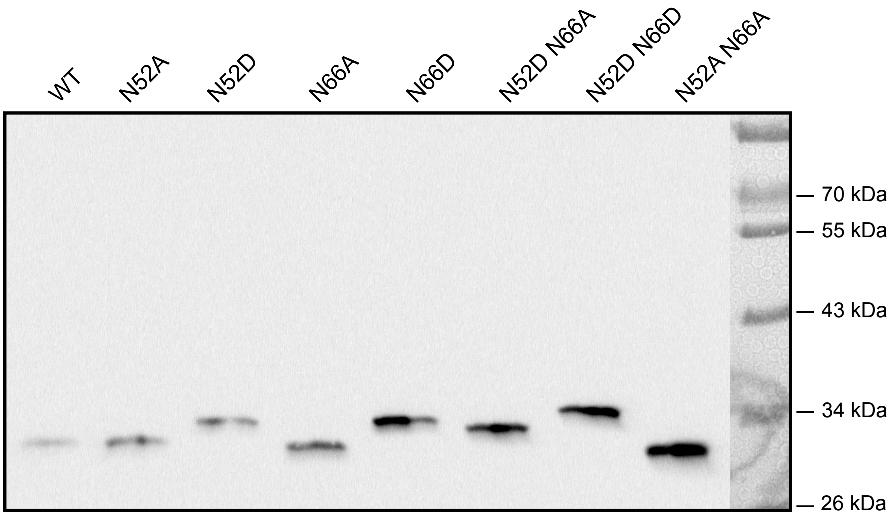


**Suppl Figure 1: Electrophoretic separation of deamido-mimics or non-deamidable mutants of Bcl-xL on Taurine-Glycine-Imidazole (TGI) gels.**

Proteins were produced by in vitro cell-free synthesis in continous exchange. 0.02% of the soluble proteins was separated on 12%TGI gels and analyzed by western blotting against Bcl-xL. Image is representative of at least 8 independent experiments.

At physiological pH, spontaneous protein deamidation converts neutral Asn to the cognate carboxylic acid, and the replacement of the -CO(NH2) group by -CO(OH) entails a 0.984 Da mass increase and a negative charge per deamidation site. These chemical changes produce a migration shift in SDS-PAGE, which is larger than expected for the ~1Da change, and further amplified on in-house TGI gels (Bobo et al., 2019; Boudier-Lemosquet et al., 2022). Distinctive migration distances can be appreciated for Bcl-xL proteoforms depending on their deamidation status. Deamido-mimic mutants include Bcl-xL with N52 and/or N66 mutated to D. Non-deamidable mutants include Bcl-xL with N52 and/or N66 mutated to A.

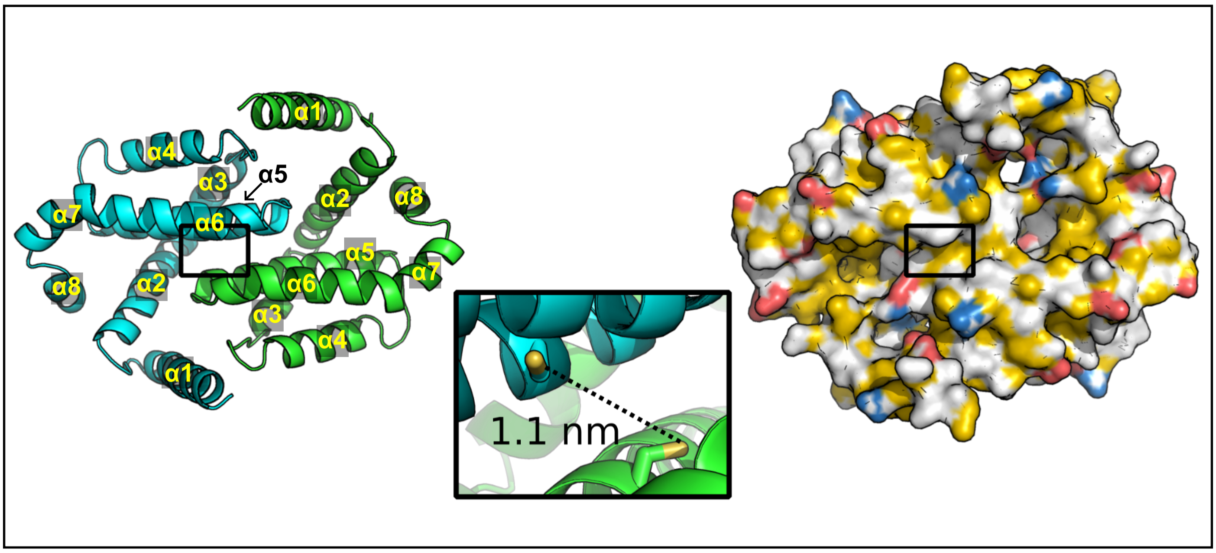


**Suppl Figure 2: Best pose of Bcl-xL non-covalent dimer**, as obtained with HADDOCK 3, using the crystal structure (PDB code: 3FDL) of Bcl-xLΔLoopΔα9 as the starting structure (see Methods). On the left, a cartoon representation of the dimer, in which one chain is colored in cyan and one in green; on the right, the electrostatic surface of the dimer, obtained according to the procedure developed by Hagemans and coworkers (Hagemans D, van Belzen IAEM, Morán Luengo T and Rüdiger SGD (2015) A script to highlight hydrophobicity and charge on protein surfaces. Front. Mol. Biosci. 2:56. doi: 10.3389/fmolb.2015.00056), and colored such that the hydrophobic regions are in yellow, the negative in red, and the positive in blue. The black square on both dimer representations indicates the position of the two Cys151 residues, drawn as sticks in the central zoom inset.

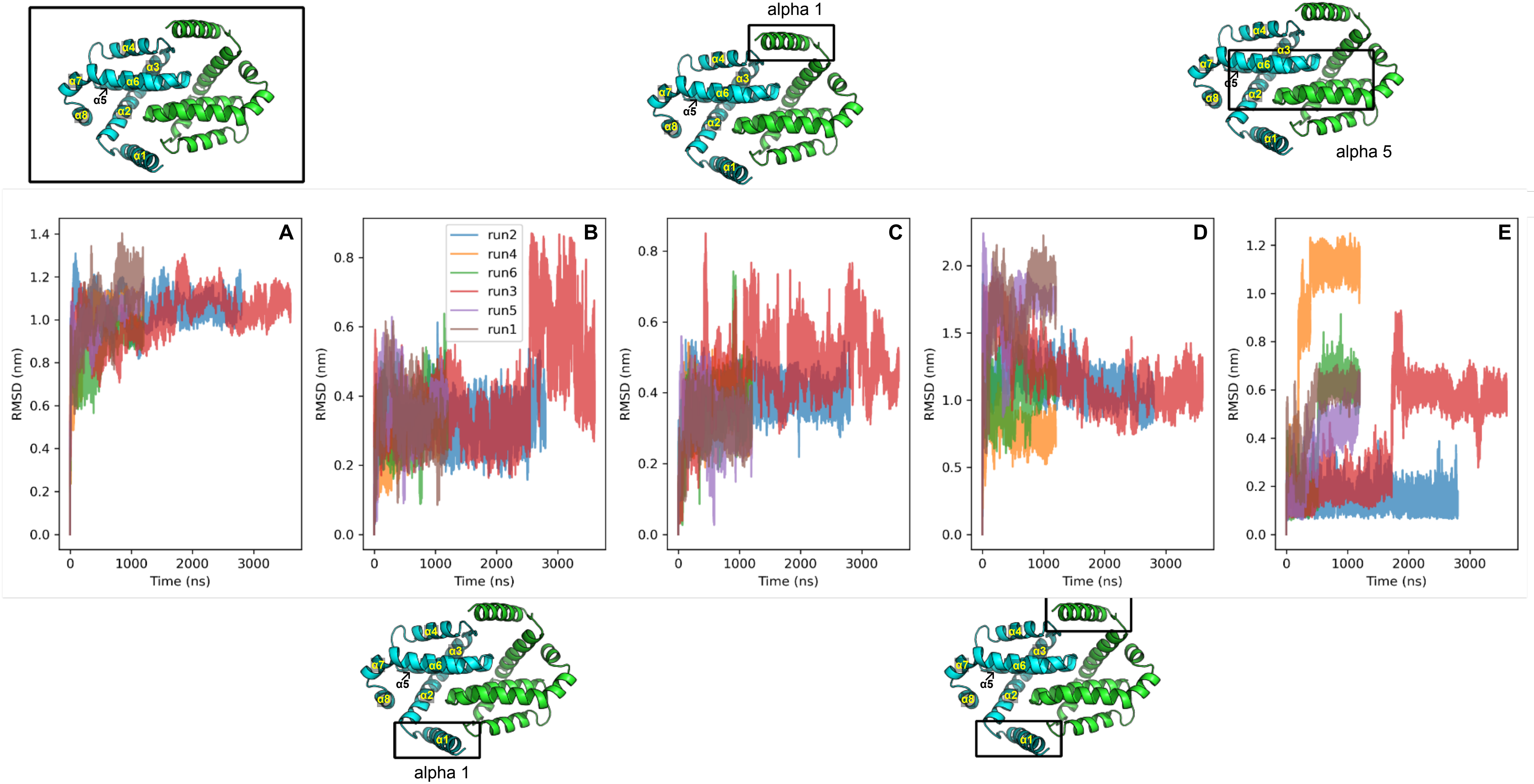


**Suppl Figure 3: Structural-dynamics analysis of Bcl-xL non-covalent dimer.** Simulations start from the best docking pose of the non-covlent dimer, and RMSD fluctuations are reported over the indicated simulation times (listed in Table 1). In the proximity of each graph, the black square indicates the analyzed region. Note that Y axis are not set to the same scale. **A-** RMSD analysis of the whole dimer shows comparable behaviors in each replica, reaching a similar plateau, suggestive of a stable structure. **B- C-** Atomic deviation of helix1 in one monomer (B) or the other (C) relative to the whole dimer or **D-** relative to one another: RMSD fluctuations suggest that helices1 are the most mobile regions of the structure, both in terms of position and orientation. **E-** Analysis of the structural dynamics of the central α5-α6 hairpin of one monomer relative to the other.

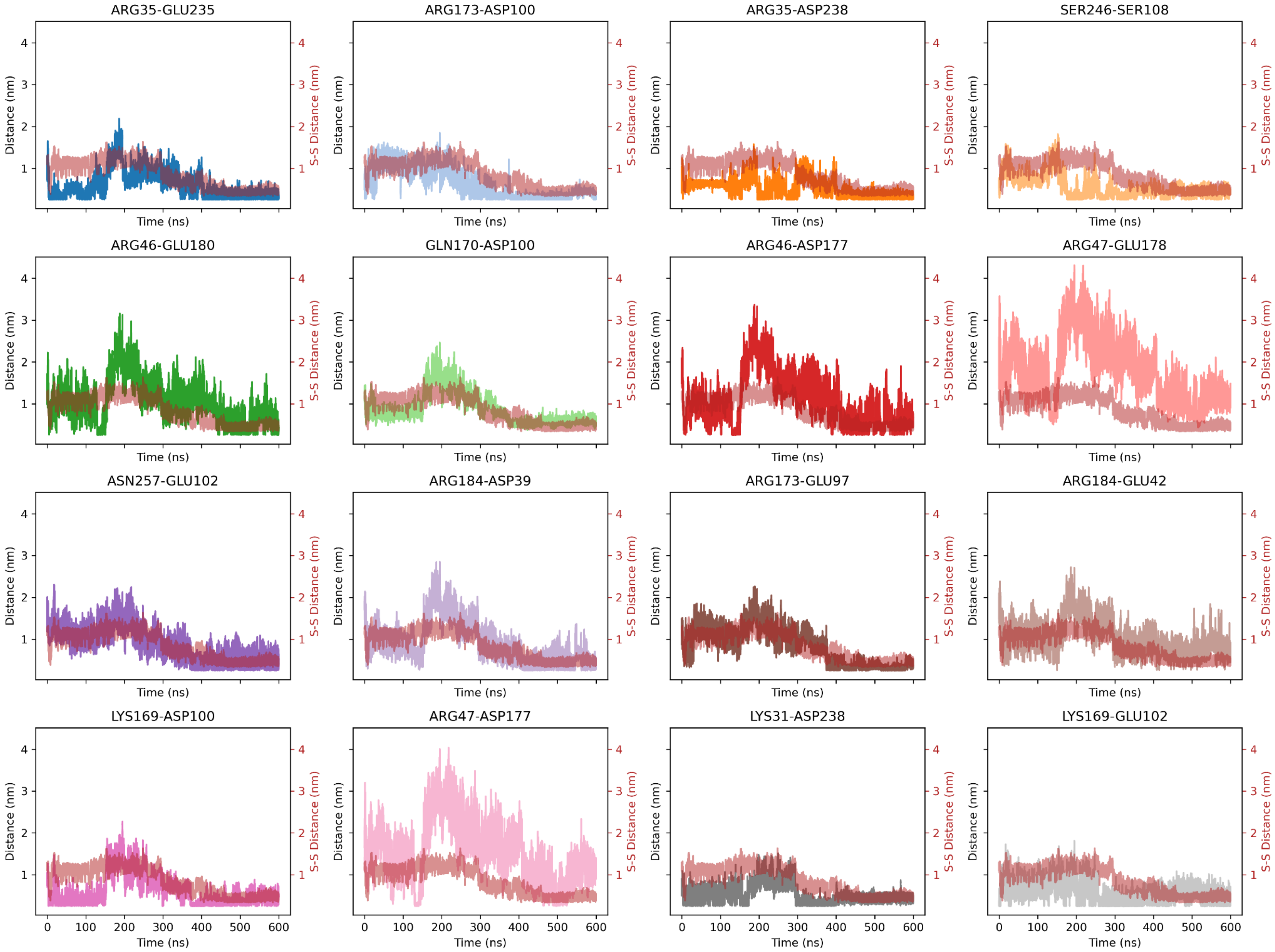


**Suppl Figure 4:** Analysis of salt bridges and hydrogen bonds between selected residues shows a marked decrease in inter-residue distances. Residues on one monomer are paired with a residue on the other monomer. For direct comparison, the reduction in the Cys151-Cys151 distance monitored in run1 of MD simulations of non-convalent Bcl-xL dimers is represented in red in the background of all the subplots. Data show consistency between paired-residues brought closer and directional sliding of the central hairpins (helices alpha5–alpha6) toward one another, supporting possible disulfide bridge formation.

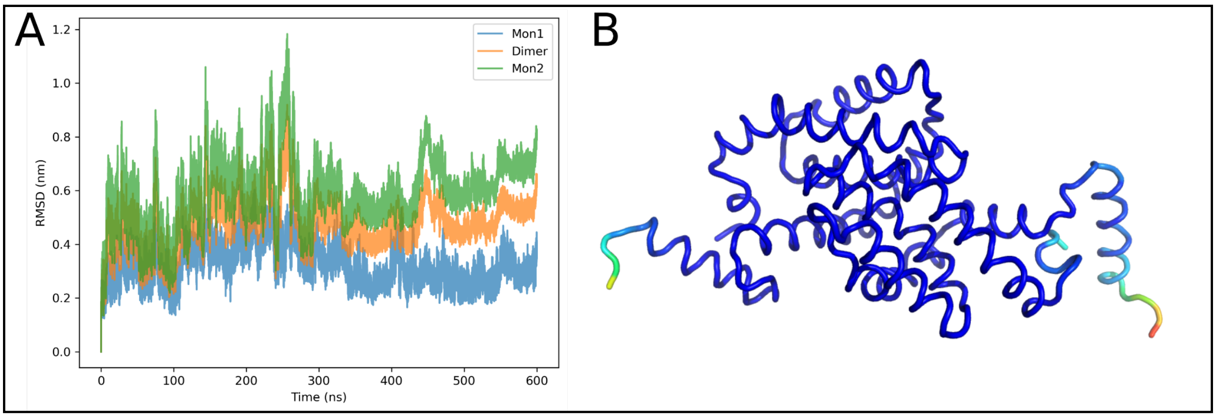


**Suppl Figure 5: C-alpha RMSD of the S-S dimer**

**A-** Using the simulation in Figure 3C, a Bcl-xL dimer was formed enforcing the formation of a C155-C151 disulfide bridge. The simulation is compatible with a stable dimer. **B-** Color-coded representation showing regions of the protein that fluctuate the most. Blue = low values ; Red = high values. The highest fluctuations are observed for helices alpha1 in each monomer.
